# Differential Contributions of Static and Time-Varying Functional Connectivity to Human Behavior

**DOI:** 10.1101/2020.07.14.203273

**Authors:** Adam Eichenbaum, Ioannis Pappas, Daniel Lurie, Jessica R. Cohen, Mark D’Esposito

## Abstract

Measures of human brain functional connectivity acquired during the resting-state track critical aspects of behavior. Recently, fluctuations in resting-state functional connectivity patterns – typically averaged across in traditional analyses – have been considered for their potential neuroscientific relevance. There exists a lack of research on the differences between traditional “static” measures of functional connectivity and newly-considered “time-varying” measures as they relate to human behavior. Using functional magnetic resonance imagining (fMRI) data collected at rest, and a battery of behavioral measures collected outside the scanner, we determined the degree to which each modality captures aspects of personality and cognitive ability. Measures of time-varying functional connectivity were derived by fitting a Hidden Markov Model. To determine behavioral relationships, static and time-varying connectivity measures were submitted separately to canonical correlation analysis. A single relationship between static functional connectivity and behavior existed, defined by measures of personality and stable behavioral features. However, two relationships were found when using time-varying measures. The first relationship was similar to the static case. The second relationship was unique, defined by measures reflecting trialwise behavioral variability. Our findings suggest that time-varying measures of functional connectivity are capable of capturing unique aspects of behavior to which static measures are insensitive.

**Author Summary:** Correlated patterns of brain activity measured in the absence of any prescribed task show meaningful temporal fluctuations. However, the manner by which such fluctuations track aspects of human behavior remains unresolved. The current report takes a data-driven approach to characterize how time-varying patterns of human brain functional connectivity differ from traditional static measures in their ability to track aspects of personality and cognitive ability. We determine that time-varying patterns of functional connectivity not only track similar aspects of behavior as do static measures, but also unique behavioral qualities as well, specifically those that reflect behavioral variability. These results highlight the importance and relevance of examining time-varying measures of functional connectivity.

## Introduction

Measuring spontaneous activity in the human brain during a task-free “resting state” has become common as this activity has been revealed to be spatially and temporally organized (Biswal et al., 1995). These patterns of resting-state functional connectivity (rsFC) are sensitive to numerous aspects of behavior measured outside the scanner, including cognitive performance (Stevens et al., 2012; Chan et al., 2014), age (Chan et al., 2014), and the extent of cognitive impairments (Alexander-Bloch et al., 2010; Rudie et al., 2013). Using rsFC data from the Human Connectome Project (Van Essen et al., 2013), a recent report utilized canonical correlation analysis (CCA) to reveal that rsFC and a myriad of behavioral measures were linked via a single mode of population covariation, thus providing a single inextricable link between stable functional brain organization and interindividual differences in behavior (Smith et al., 2015).

The majority of neuroimaging studies have investigated rsFC by assuming that it is stable across the measurement period. However, a recent emphasis has been placed on determining whether, and to what degree, rsFC systematically varies in time (Calhoun et al., 2014). While it is likely that some measurable fluctuations are due to noise or non-neural, physiological signals (Hutchison et al., 2013; Lindquist et al., 2014; Hindriks et al., 2016; Duff et al., 2018; Lurie et al., 2019), there is evidence that these rapidly evolving changes have a neuronal basis (Chang and Glover, 2010; de Pasquale et al., 2010; Brookes et al., 2014; Thompson, 2018). Moreover, analysis of time-varying FC might reveal new relationships to behavior unobtainable by static analyses (Cohen, 2018; Kucyi et al., 2018). For example, there is recent evidence that fluctuations of task-based FC track aspects of cognitive control (Khambhati et al., 2018) and attention (Sadaghiani et al., 2015), suggesting that flexible network reconfiguration indexes trial-by-trial performance.

It is important to consider ways in which static and time-varying FC differ, and how these differences impact the way each modality encodes aspects of behavior. Whereas static measures provide a snapshot of the stable organization of the brain, time-varying measures index higher-order relationships between brain regions. Such measures include the degree to which functional networks vary their interconnectivity with other networks, the change in global organizational structure, and how the global FC profile transitions between different functional substates (Vidaurre et al., 2017; Shine, Breakspear et al., 2018). Thus, it is likely that measures of static and time-varying FC encode different behavioral features, however a precise characterization of this relationship is missing. Studies have focused on either one type of connectivity (static: Smith et al., 2015, time-varying: Casorso et al., 2019), or on specific behaviors (Rosenberg et al., 2016), but only one study attempted to simultaneously disentangle static and time-varying FC’s behavioral relevance (Liegeois et al., 2019). In that study, the authors report that measures of time-varying FC tracked both task-based behavior and self-reported personality traits, whereas static measures only captured self-reported traits. Although it took advantage of a powerful sample size, this study only had access to basic measures of human behavior, lacking access to measures typically employed by cognitive neuroscientists studying working memory, cognitive control, and executive function.

In order to directly address the behavioral differences captured by static and time-varying measures of FC, we utilized a dataset of resting-state blood-oxygen-level-dependent (BOLD) data collected alongside a battery of complex behavior and personality measures. These measures ranged across working memory, executive functioning, processing speed, affect, and impulsivity. Building off Smith and colleagues (2015), we leveraged CCA to determine whether there exist modes of covariation between behavior and static, as compared to time-varying, rsFC. Static rsFC was estimated by computing a node-node correlation matrix across all regions of the brain. Time-varying rsFC was estimated by fitting a Hidden Markov Model (HMM) to the data. The HMM allowed for the characterization of, and transition likelihood between, multiple latent “states” in a data-driven fashion as fast as the modality allowed, overcoming limitations imposed by classical sliding-window methods (Hutchison et al., 2013). The HMM has been used to characterize brain dynamics across multiple neuroimaging modalities during rest (Baker et al., 2014; Vidaurre et al., 2017) and task (Vidaurre et al., 2018).

Using static FC, CCA revealed a single relationship primarily defined by variance in measures of personality and affect, as well as task-general behavioral features. With time-varying FC, CCA instead revealed two (orthogonal) relationships. The first was highly similar to that found using static FC. However, the second was specific to time-varying FC, and was defined by variance in trialwise measures of reaction time to processing speed and working memory tasks, as well as measures tapping into overall processing accuracy. These results suggest that there exists meaningful information in the temporal fluctuations of rsFC patterns which can explain aspects of human behavior to which previous methods of analysis have been insensitive.

## Methods

### Participants

Twenty-three healthy young adult participants (mean age = 28.26 years, SD = 4.52 years, 10 females) were recruited for a repeated measures study to participate in two or three sessions. Five participants were unable to attend the third study session as a result of having moved away from the state of California. As a result, only 18 participants were included in the third session (mean age = 27.67 years, SD = 4.64 years, 8 females). All participants were native English speakers, had normal or corrected-to-normal vision, and had normal hearing. Participants were excluded for any history of neurological or psychiatric disorders, use of psychotropic drugs, a history of substance abuse, or MRI contraindications. All participants provided written informed consent according to the procedures of the UC Berkeley Institutional Review Board.

### Experimental Design and Procedure

Participants underwent one practice session approximately one week (mean = 6 days, SD = 2.37 days) before their first testing session. They then completed two or three identical testing sessions. Testing sessions 1 and 2 were separated by approximately one week (mean = 8 days, SD = 1.47 days), while testing sessions 2 and 3 were separated by approximately one year (mean = 399 days, SD = 28.73 days).

Each session began with two 6-minute resting-state scans in the MRI machine, in which participants were instructed to stay awake with their eyes open and fixate on a crosshair. During the first session, the resting-state scans were followed by a structural scan. Immediately after the MRI scan, participants completed two self-report questionnaires and a task outside of the scanner: a Visual Analogue Scale (VAS, McCormack et al., 1988), the Barratt Impulsiveness Scale (BIS, Patton et al., 1995) and a Box Completion Task (Salthouse, 1996). Immediately following completion of the questionnaires and task, participants then completed four computerized cognitive tasks in counterbalanced order (different orders across participants and for each session): a Stroop task (Stroop, 1935), a digit symbol substitution task (DSST; Rypma et al., 2006), a spatial working memory (WM) task (Kuo et al., 2011), and a color WM task (Zhang & Luck, 2008).

The BIS is a survey that determines measures of impulsivity along a set of three sub-traits: “Attentional,” “Motor,” and “Non-planning”. The VAS has participants make a mark along a line segment in which one side represents “Not” and the other side “Extremely” for the following items: “Anxious,” “Happy,” “Sad,” “Nauseous,” “Drowsy,” “Jittery,” “Fatigued,” and “Dizzy”. Participant responses are measured as the distance (in centimeters) away from the “Not” end of the line. The Box Completion Task requires that participants use a pencil to fill in the fourth side of an open-ended square as rapidly as possible. The measure of interest is the duration of time it takes to complete 100 squares.

In the Stroop task, color words (blue, red, green, yellow) or animal names (horse, bird, cat, dog) printed in different colors (blue, red, green, yellow) were presented on the left side of the computer screen. Participants had to indicate the font color by pressing one of four buttons. For ease of task performance color-to-button mappings were presented at the bottom part of the screen throughout the duration of the experiment. Participants used the four fingers of their right hand for responding with color-to-button mappings randomly assigned to participants. Compatible, neutral, and incompatible trials were presented with equal probability. In compatible trials, color and word were the same. In neutral trials, the task-irrelevant dimension (e.g., word meaning) was not related to the task (e.g., animal names). In incompatible trials, color and word differed. Each Stroop session was ten minutes long and comprised 8 blocks of 36 trials each. The stimuli were presented for 300 ms with an interstimulus interval of 1700 ms. The measures of interest included the difference score, in milliseconds, between the median response time of correct responses to trials in which there was an incongruity between the word and color (incongruent trials: i.e., the word “RED” in blue text) and the median response time of correct responses to a trial in which the color of the text matched the word (congruent trials: i.e., the word “RED” in red text). Moreover, we also focused on the standard deviation of this response time difference, as well as the accuracy on incongruent trials. We chose not to compute a difference score for accuracy as individual differences for accuracy on congruent trials was likely to be minimal.

The DSST required that participants indicated via button press whether a presented symbol-number pair correctly matched an on-screen answer key. Nine symbols were paired with numbers 1 through 9 and the answer key was shown at the top of the screen on every trial. 140 pairs were presented in which the symbol-number pair either matched (50%), or did not match (50%), the provided answer key. Pairs were presented on screen for 4000 ms, during which the participant could indicate their response. Participants were instructed to respond as rapidly and as accurately as possible. Measures of interest included the overall accuracy, median reaction time, and standard deviation of reaction time, for match and non-match trials separately.

The spatial WM task (“Spatial WM”) required that participants initially encode and retain the color of a rapidly presented set of colored squares. The task followed a 2 (load: 2 vs. 4) × 2 (cue onset: early vs. late) design. Participants viewed an array of 2 or 4 colored squares for 180 ms prior to retaining this information over a 900 ms delay period. In the early-cue condition, a cue appeared in the location of where one of the squares had previously been after 200 ms (and stayed on screen for the remaining 700 ms). In the late-cue condition, the cue appeared after 800 ms (and stayed on screen for the remaining 100 ms). Next, participants had to indicate whether a newly presented colored square, among an array of 2 or 4 colored squares, matched the color of the spatially cued square prior to the delay. The new array remained on screen for 1920 ms. Participants were instructed to respond as accurately and as quickly as possible. In total, participants completed 240 trials, with 60 trials coming from each condition. Measures of interest included percent accuracy, median reaction time, and the standard deviation of reaction time, across both cognitive loads, for match and non-match trials separately.

The color working memory task (“Color WM”) required that participants initially encode the colors of 3 squares rapidly presented on screen for 1000 ms. Following a delay of 500 ms, a visual cue to the location of one of the squares appeared for 500 ms. After a 1250 ms delay, a distractor color appeared on screen for 500 ms. Following another delay of 1250 ms, the participants were then presented with a colorwheel for 3000 ms and were instructed to move the cursor along the wheel in a continuous fashion until the selected color matched the color of the cued square being held in memory. Participants completed 40 trials in total and were provided a 5 second break after the end of the 20th trial. Measures of interest included the median and standard deviation of reaction time and error angle (calculated as the difference in degrees along the colorwheel between the correct answer and the response provided by the participant) across all responses.

During the practice session, participants completed the four cognitive tasks so as to familiarize themselves with the tasks before the testing sessions. The purpose of this session was to minimize practice effects. The testing sessions were all identical. The final testing session was conducted on the same MRI machine as the previous sessions, but in a different location. Reliability tests ensured that MRI effects (such as signal-to-noise ratio and artifacts) were not different across the two locations.

### Factor Analysis of the Behavioral Data

All 31 behavioral measures were included in the analyses and subjected to a factor analysis. Six measures each came from the Spatial WM task and the DSST: percent accuracy, median reaction time, and the standard deviation of reaction times for match and non-match trials. Three measures came from the Stroop task: percent accuracy on incongruent trials, median reaction time difference between congruent and incongruent trials, and the standard deviation of the reaction time difference between congruent and incongruent trials. Four measures came from the Color WM task: median and standard deviation of response error, as well as median and standard deviation of reaction times. All eight measures from the VAS were included, as well as the scores of the three sub-traits of the BIS. Lastly, the time to complete all 100 squares for the Box Completion Task was included.

We clustered the behavioral data into 8 factors using MATLAB’s *factoran* function and allowed for promax oblique rotation (Supplementary Figure S1). We labeled these factors qualitatively by observing which behavioral measures loaded highest on each factor. We chose 8 factors as it most cleanly separated tasks from one another and grouped together correlated measures.

### fMRI Data Acquisition

Imaging data were collected on a 3-Tesla Siemens MAGNETOM Trio whole-body MR scanner using a 12-channel head coil at the University of California, Berkeley Henry H. Wheeler Jr. Brain Imaging Center. Whole-brain functional data were acquired in two runs using a T2*-weighted echo-planar imaging (EPI) pulse sequence (180 volumes/run, 37 interleaved axial slices parallel to the AC-PC line, slice thickness 3.5 mm, interslice distance = 0.7 mm, TR = 2000 ms, TE = 24 ms, FA = 60°, matrix 64 × 64, field of view 224 mm). A high-resolution T1-weighted structural 3D MP-RAGE was also acquired (160 slices, slice thickness 1 mm, TR = 2300 ms, TE = 2.98 ms, FA = 9°, matrix 256 × 256, field of view 256 mm). An LCD projector back-projected a fixation cross for the resting-state scan onto a screen mounted to the RF coil.

### fMRI Data Processing

Preprocessing of the imaging data were performed using fMRIPrep 1.1.4 (Esteban, Markiewicz et al., 2018; Esteban, Blair et al., 2018), which is based on Nipype 1.1.1 (Gorgolewski et al., 2011). The T1-weighted (T1w) image was corrected for intensity non-uniformity (INU) using N4BiasFieldCorrection (ANTs 2.2.0, Tustison et al., 2010), and used as T1w-reference throughout the workflow. The T1w-reference was then skull-stripped using ANTs BrainExtraction (ANTs 2.2.0), using OASIS as target template. Brain surfaces were reconstructed using recon-all (FreeSurfer 6.0.1, Dale, Fischl, & Sereno, 1999), and the brain mask estimated previously was refined with a custom variation of the method to reconcile ANTs-derived and FreeSurfer-derived segmentations of the cortical gray-matter of Mindboggle (Klein et al., 2017). Spatial normalization to the ICBM 152 Nonlinear Asymmetrical template version 2009c (MNI152NLin2009cAsym, Fonov et al., 2009) was performed through nonlinear registration with ANTs Registration (ANTs 2.2.0, Avants et al., 2008), using brain-extracted versions of both T1w volume and template. Brain tissue segmentation of cerebrospinal fluid (CSF), white-matter (WM) and gray-matter (GM) was performed on the brain-extracted T1w using fast (FSL 5.0.9, Zhang, Brady, & Smith, 2001).

For each of the BOLD runs found per participant (across all sessions), the following preprocessing was performed. First, a reference volume and its skull-stripped version were generated using a custom methodology of fMRIPrep. Head-motion parameters with respect to the BOLD reference (transformation matrices, and six corresponding rotation and translation parameters) were estimated before any spatiotemporal filtering using mcflirt (FSL 5.0.9, Jenkinson et al., 2002). BOLD runs were slice-time corrected using 3dTshift from AFNI. The BOLD time series (including slice-timing correction when applied) were resampled onto their original, native space by applying a single, composite transform to correct for head-motion and susceptibility distortions. These resampled BOLD time series will be referred to as preprocessed BOLD in original space, or just preprocessed BOLD. The BOLD reference was then co-registered to the T1w reference using bbregister (FreeSurfer) which implements boundary-based registration (Greve & Fischl, 2009). Co-registration was configured with nine degrees of freedom to account for distortions remaining in the BOLD reference. The BOLD time series were resampled to MNI152NLin2009cAsym standard space, generating a preprocessed BOLD run in MNI152NLin2009cAsym space. Several confounding time series were calculated based on the preprocessed BOLD: framewise displacement (FD), DVARS, and three region-wise global signals. FD and DVARS were calculated for each functional run, both using their implementations in Nipype (following the definitions by Power et al., 2014). The three global signals were extracted within the CSF, the WM, and the whole-brain masks (i.e., global signal). The head-motion estimates calculated in the correction step were also placed within the corresponding confounds file. All resamplings were performed with a single interpolation step by composing all the pertinent transformations (i.e. head-motion transform matrices, susceptibility distortion correction when available, and co-registrations to anatomical and template spaces). Gridded (volumetric) resamplings were performed using ANTs ApplyTransforms, configured with Lanczos interpolation to minimize the smoothing effects of other kernels (Lanczos, 1964).

Further post-processing included removal of artifactual signals from the time series data. We regressed out the six head-motion estimates, the mean white matter signal, the mean CSF signal, and both the derivatives of these time series and the quadratic expansions of both the original artifactual time series and the derivative time series. In addition, a bandpass filter from 0.01 to 0.1 Hz was applied to the data.

### Static Functional Connectivity

In order to obtain measures of FC, we first measured the mean BOLD signal across all voxels contained within each node of our brain atlases. Cortical nodes were taken from the 400-node Local-Global atlas (Schaeffer et al., 2018). 21 subcortical nodes were taken from the Harvard-Oxford atlas (Makris et al., 2006). 22 cerebellar nodes were taken from the AAL atlas (Tzourio-Mazoyer et al., 2002). Four cortical nodes in bilateral anterior temporal pole regions had to be removed from all analyses due to insufficient coverage (less than 25% of voxels contained data) in one or more participants in one or more scans. This left data from 439 nodes distributed across the entire brain.

Scans were concatenated within session, per participant, in order to increase reliability of the measured FC profile for each session. To remove spurious data differences between sessions, each session’s data was standardized. FC was measured as the Pearson correlation coefficient between every node and all other nodes for which there was sufficient coverage.

### Hidden Markov Model

#### Setup

The HMM derives brain dynamics based on BOLD time series parcellation data. The HMM assumes that the time series data are characterized by a number of states that the brain cycles through at different times throughout the scanning period (Baker et al., 2014).

At each time point *t* of brain activity, the observed time series data was modeled as a mixture of multivariate Gaussian distributions. Each one of these Gaussian distributions corresponded to a different state *k* and was described by first-order and second-order statistics (activity [*μ*_*k*_] and FC [*Σ*_*k*_], respectively) that can be interpreted as the activity and FC of each state. Using notation, if *x*_*t*_ describes the BOLD data at each time point *t*, then the probability of being in state *k* is assumed to follow a multivariate Gaussian distribution

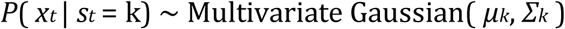

In turn, we modeled how transitions between states took place. The basic Markovian principle that describes the transition between states assumes that the probability of the data being in state *k* at time *t* relates only to the probability of being in state *l* at time *t-1*. This can be described by the following equation

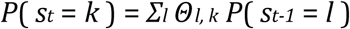

where Θ_*l, k*_ is the transition probability from state *l* to state *k*. Taken together, the HMM infers the *P*(*s*_*t*_ = *k*) probabilities for each state *k* and time *t* (state time courses) as well as the transition probabilities Θ_*l, k*_ and the statistics of each state (*μ*_*k*_, *Σ*_*k*_) that best describe the data. To make inference tractable, a variational Bayes algorithm was used that works by minimizing the Kullback-Leibler divergence between the real and the modeled data (Wainright & Jordan, 2007).

The input time series data for the HMM was the total time series data for all participants and all sessions (for the last session there were only 18 participants). Specifically, across the three sessions and for all participants we concatenated the processed functional time series and obtained a matrix of dimensions: (180×23 + 180×23 + 180×18) × number of regions of interest (439) (Vidaurre et al., 2017). Data were standardized for each participant prior to running the HMM. To control the dimensionality in the final data matrix, a principal component analysis (PCA) dimensionality-reduction technique was applied on the concatenated time courses using 25 components. Finally, the number of states for the HMM was chosen as 12. Both of these settings were similar to the previous work that introduced the use of the HMM on fMRI data (Vidaurre et al., 2017).

#### Inference

Running the HMM with these parameters resulted in a data matrix of dimensions (# time points x # participants) x # states. Each row represented the probability of each state being active at each timepoint for each participant. Additional quantities related to the temporal characteristics of each state could then be obtained. First, we quantified the proportion of time that an individual resided in the state during the scan acquisition (fractional occupancy; FO). Additionally, the switching rate was defined as the difference between the probability of activating a state at time *t* and activating a state at time *t+1* summed over all states and over all time points and divided by the number of time points. The HMM also provided each state’s mean activity and connectivity *μ*_*k*_ and *Σ*_*k*_, respectively. Finally, the HMM also provided the state transition probability matrix of dimensions (# states x # states) where each matrix entry (*k, l*) quantified the transition probability of going from state *k* to state *l*.

An agglomerative hierarchical clustering algorithm was applied to the transition probability matrix in order to determine whether there existed a temporal structure in the data, as had previously been shown with resting-state FC data from the Human Connectome Project (Vidaurre et al., 2017). This analysis starts by classifying each data point as a separate cluster and progressively combines clusters of data at different hierarchical levels: similar data are clustered at a low level of hierarchy and less similar data are clustered at a higher level of hierarchy (Hastie et al., 2009). We used the *linkage* function as implemented in MATLAB with default settings (method= ‘single’, distance= ‘euclidean’). We regarded each identified cluster as one metastate. In turn, the metastate time courses were considered as the sum of the time courses of the individual states that comprised them. Fractional occupancy and switching rate of the metastates were calculated as in the case for single states.

### Spatial Characterization of States

In order to spatially characterize the derived states, we thresholded the activity maps of each state to include the top 40% of both positive and negative activations. We then spatially overlapped each state with the 10 resting-state networks described in Smith et al. (2009) to obtain an overlap index for each network. The index was calculated by counting the number of voxels that were included in the thresholded map and then dividing these by the size of the resting-state network under consideration in order to account for size bias.

### Canonical Correlation Analysis (CCA)

To relate the behavioral measures to static and time-varying FC we used CCA (Figure 1). CCA finds correlations between multidimensional data wherein potential relationships may be present (Hotelling, 1936). This is a more principled approach compared to conducting all potential correlations and correcting for multiple comparisons. Specifically, this analysis finds maximal correlations between two sets of variables, X (n x d1) and Y (n x d2), where d1 and d2 are the number of variables used in X and Y respectively, and n is the number of observations for each variable. It produces two matrices, A and B, such that the variables U=AX and V=BY are maximally related. CCA values were obtained from the MATLAB *canoncorr* function. It is worth noting that like the PCA, this function can produce more than one mode, with each mode ranked by the covariance that can be explained between X and Y.

**Figure 1.**
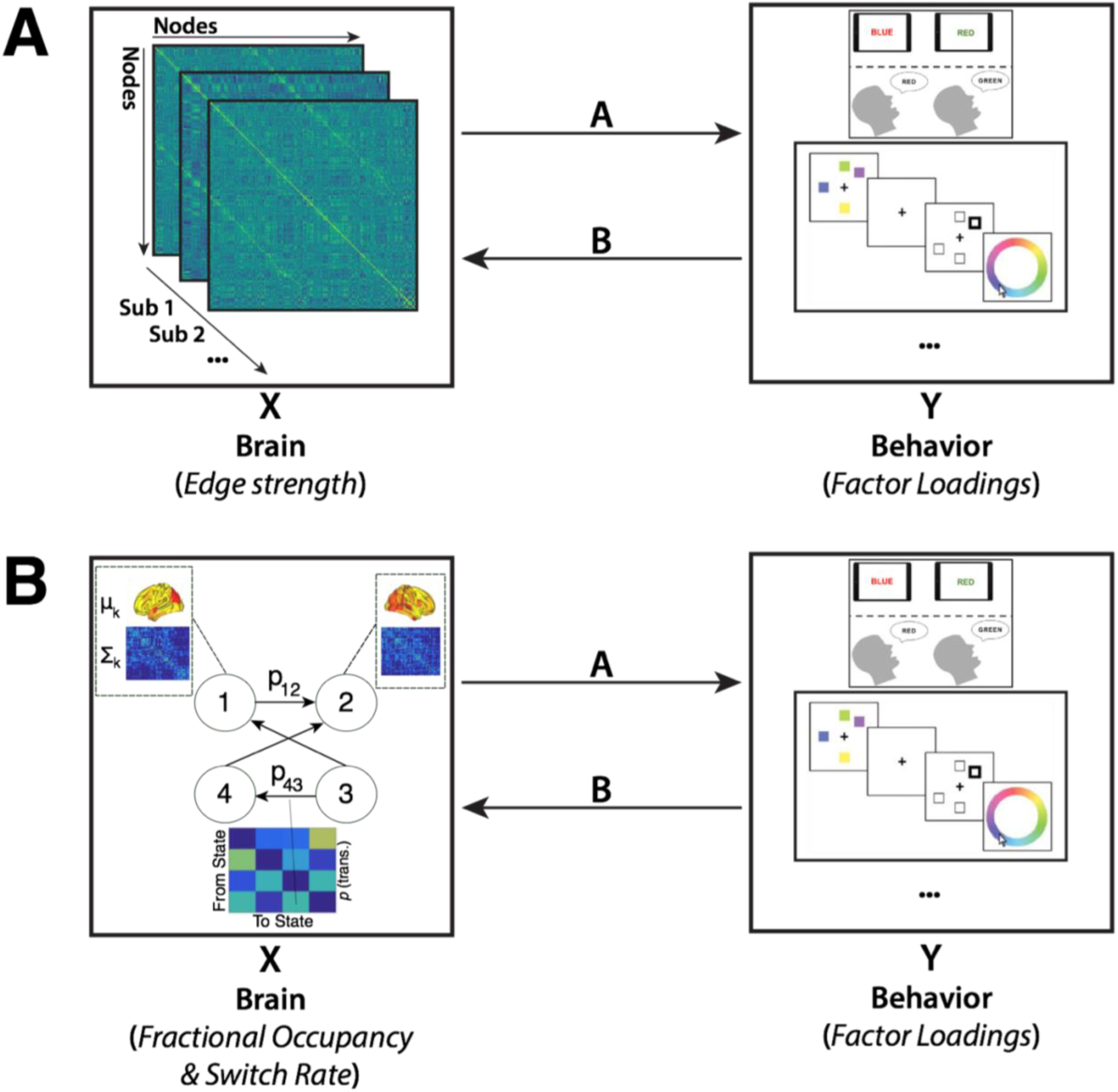
Methodology Overview. CCA was performed on two different datasets, which were matched for measures of behavior but differed with regard to the rsFC data included. The first CCA (A) included measures of static FC (i.e., the node-to-node connectivity strength), while the second CCA (B) included measures of time-varying FC. Measures of time-varying FC were derived by fitting a Hidden Markov Model to the BOLD time series.

We conducted two separate CCAs. First, we designated the factors of the behavioral data as Y, and the edgewise static FC strength as X (n = 96,141). In a second CCA, Y remained the same, but we varied X. Specifically, we designated X as the fractional occupancy of each HMM state (n = 12) and temporally defined metastate (n = 3), as well as the mean switching rate across states (n = 1) and metastates (n = 1) separately. As a final preprocessing step, the dimensionality of the static FC data was reduced using PCA as described in Smith et al. (2015), retaining the top 13 components. No dimensionality reduction was required for the HMM data as the number of variables was low. However, we performed an analysis of the HMM data using PCA and report the results in the Supplementary Materials (Supplementary Figure S3) where it can be seen that the results are highly similar to the case when PCA is not employed.

Statistical significance of the CCA analyses was estimated as follows. We calculated 10,000 permutations of the rows of X relative to Y, respecting the within-participant structure of the data, and recalculated the CCA mode for each permutation in order to build a distribution of canonical variate pair correlation values (i.e. <U,V>). By comparing the outcome from the CCA of the true data to the shuffled data, we found that each mode of covariation discovered with the true data was highly significant (p < 1/10,000). In addition, a cross-validation approach was adopted in order to assess the robustness of the discovered mode(s) (as described in Smith et al., 2015). Across 1,000 runs, we ran CCA on a randomly selected set of 80% of the data, respecting the within-participant nature of the data, and stored the resultant U and V. We then estimated the mode on the held-out 20% of data and determined the significance of the estimated mode employing the same permutation significance testing procedure as before. These estimated modes were found to be highly significant, with the correlation between the derived canonical weight vectors in the test dataset being very robust (replicating the results from Smith et al., 2015).

Post-hoc correlations of the values of X (Y-respective) with the columns of the significant mode U (V-respective) were used to quantify the contributions (positively or negatively) of each behavioral measure with the CCA mode. In other words, we quantified the extent to which the Y variables were loaded/weighted on the CCA mode. There is no clear cutoff at which one finds a significant correlation value and thus correlation values are reported in isolation.

### Validation of CCA Analysis

We validated the identified CCA modes by comparing outcomes across a range of behavioral factors (behavior) and FC principal components (static FC, time-varying FC). The number of behavioral factors ranged from 1 to 9, while the number of static FC principal components ranged from 1 to 20 and the number of time-varying FC principal components ranged from 1 to 17 (see Supplementary Materials). For the static case, we ran CCA on each combination and stored the resulting post-hoc correlations for each behavioral measure (i.e., with respect to FC), and computed the Pearson correlation between these values across all 200 combinations (Supplementary Figures S2, S3). For combinations that included two or more behavioral factors, we found that the discovered canonical covariate modes were highly similar, with Pearson correlations tending to be very highly positive (i.e., greater than r = 0.90) as well as very highly negative (i.e., less than r = −0.90). This bimodal distribution at the extremes of the correlation range indicates that the discovered modes were highly preserved in structure (i.e., the same behavioral measures loaded highly). We determined the optimal combination (i.e., 8 behavioral factors, 13 FC PCs) by selecting either (A) the most significant canonical covariate pair (i.e. U x V), or (B) in cases where multiple pairs had the same maximal 1 / 10,000 permutation significance value, determining if combinations were highly similar after a certain number of factors or components were included, and then taking the smallest number of factors and components that produced this outcome, restricted by those that had a significant permutation value.

## Results

### Factor Analysis

Brain and behavioral data were obtained as described in the Methods. We used factor analysis to reduce the 31 behavioral measures to 8 factors (Supplementary Figure S1). The first factor, referred to as “Processing Reaction Time”, had DSST median and standard deviation reaction time measures for both match and non-match trials loading highly positively. The second factor was referred to as “Task General” because it contained a mixture of measures across multiple tasks, with positive loadings from the Spatial WM task percent accuracy (match trials), the Stroop task percent accuracy, and “Anxious” on the VAS, and negative loadings on the Stroop task median reaction time and the DSST percent accuracy (both match and non-match trials). The third factor, referred to as “Working Memory Reaction Time,” had the Spatial WM task median and standard deviation reaction time measures, for both match and non-match trials, loading highly positively. The fourth factor, referred to as “Working Memory Precision Reaction Time,” had two Color WM task measures loading highly positively: median and standard deviation of reaction time. The fifth factor, referred to as “Affect,” had the VAS measures “Sad” and “Happy” loading highly positively and negatively, respectively. The sixth factor, referred to as “Processing Accuracy,” had only the DSST percent accuracy on match trials loading highly positively. The seventh factor, referred to as “Arousal,” had high positive loadings for both the “Drowsy” and “Jittery” VAS measures. Finally, the eighth factor, referred to as “Impulsivity,” included high positive loadings of all three BIS measures.

The first (“Processing Reaction Time”), third (“Working Memory Reaction Time”), and fourth (“Working Memory Precision Reaction Time”) factors all contain measures of both the median and standard deviation of reaction time across the DSST, Spatial WM, and Color WM tasks, respectively, and therefore reflect aspects of within-task stability (median reaction time) and within-task variability (standard deviation of reaction time). In contrast, the second (“Task General”) and sixth (“Processing Accuracy”) factors only contain task measures of accuracy and/or median reaction time, and thus only reflect aspects of within-task stability. Lastly, the fifth (“Affect”), seventh (“Arousal”), and eighth (“Impulsivity”) factors all contain measures that reflect the personality and mood of the participant.

### Canonical Correlation Analysis: Static Functional Connectivity

CCA was used to find a mode of population covariation between behavior and static FC. The CCA included the behavioral data in 8-factor space, as well the static rsFC data in 13-principal component space, based on the validation we performed (see the “Validation of CCA Analysis” section of the Methods for details). The CCA revealed a single mode of covariation between these two datasets (Figure 2). To assess the statistical significance of the discovered modes of covariation, we followed the permutation and cross-validation procedure as outlined in Smith and colleagues (2015; see “Canonical Correlation Analysis (CCA)” section in Methods, and Figure 2 A, B).

**Figure 2.**
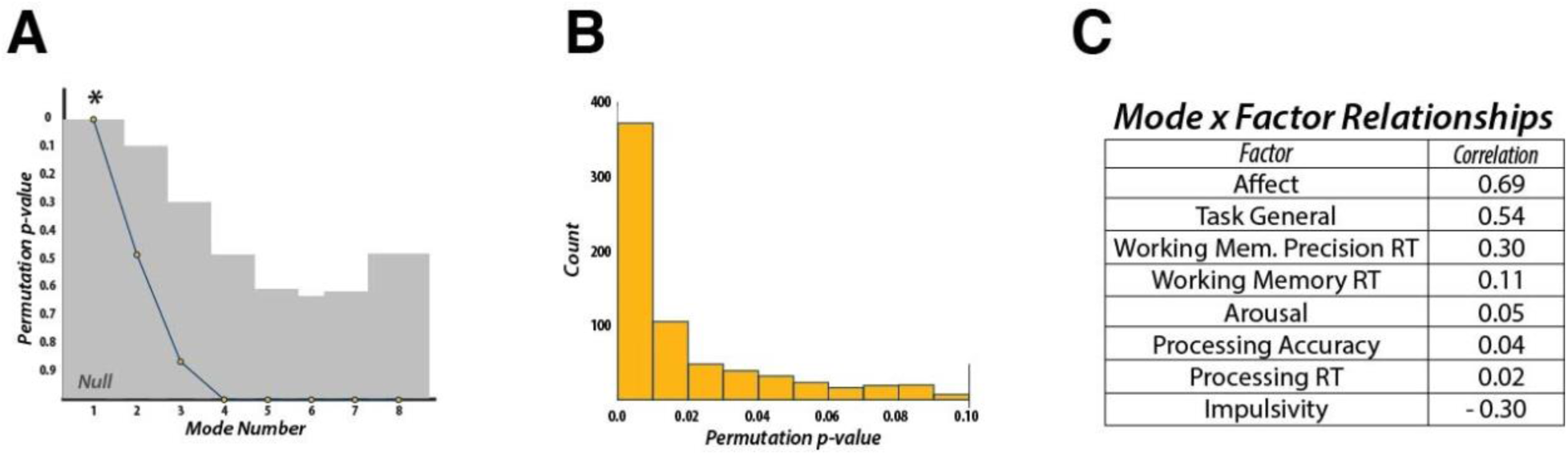
Canonical Correlation Analysis – Static Functional Connectivity. (A) CCA can discover as many modes of covariation as the lowest rank of each dataset (i.e., 8 behavioral factors). Statistical significance was found only for the first discovered mode. (B) Additional cross-validation of the discovered mode revealed that the first mode was statistically robust across the majority of the 1,000 folds. (C) Post-hoc correlations for the discovered mode and the 8 behavioral factors revealed that measures of “Affect” and “Impulsivity”, as well as a “Task General” factor, dictated the structure of the mode. RT: Reaction Time.

We used post-hoc correlations between the discovered mode and the behavioral factors to evaluate the contribution of each factor to the mode, with respect to the static FC data. This mode was defined by highly positive weights for the “Affect” (r = 0.69), “Task General” (r = 0.54), and “Working Memory Precision Reaction Time” (r = 0.30) factors, and a highly negative weight for the “Impulsivity” (r = −0.30) factor. All other factors had correlation values below an absolute value of 0.11. These results indicate that static connectivity might encode more general behavioral and personality features rather than information that may relate more to task, or trial-specific, behavior.

### Canonical Correlation Analysis: Time-Varying Functional Connectivity

We next assessed whether any relationships existed between time-varying FC and behavior. To quantify the time-varying FC profile in each participant we fit the resting-state BOLD data with a HMM. This model works by finding relevant states and their associated spatial (activity, connectivity) and temporal (fractional occupancy, switching rate) characteristics (see the “Hidden Markov Model” section in the Methods). After fitting the HMM, we identified 12 states that were representative of brain dynamics across all participants (Figure 3). Previous work has shown that the transition probabilities between HMM states derived from resting-state data is structured (Vidaurre et al., 2017). Specifically, there are certain sets of states, or “metastates”, that are more temporally coherent than others. In other words, if a participant visits a state within one metastate they are more likely to stay within that metastate compared to transitioning to another metastate. Hierarchically clustering the transition probability matrix resulted in three main clusters. One included two states, another included nine states, and the third included a single state. These results are similar to those found previously with the Human Connectome Project dataset (Vidaurre et al., 2017), indicating that even with our comparatively small sample size, we could reliably estimate brain dynamics. For completeness, we included all twelve states in our analysis; however, our results remained unchanged when we excluded the state that failed to cluster with the other states.

**Figure 3.**
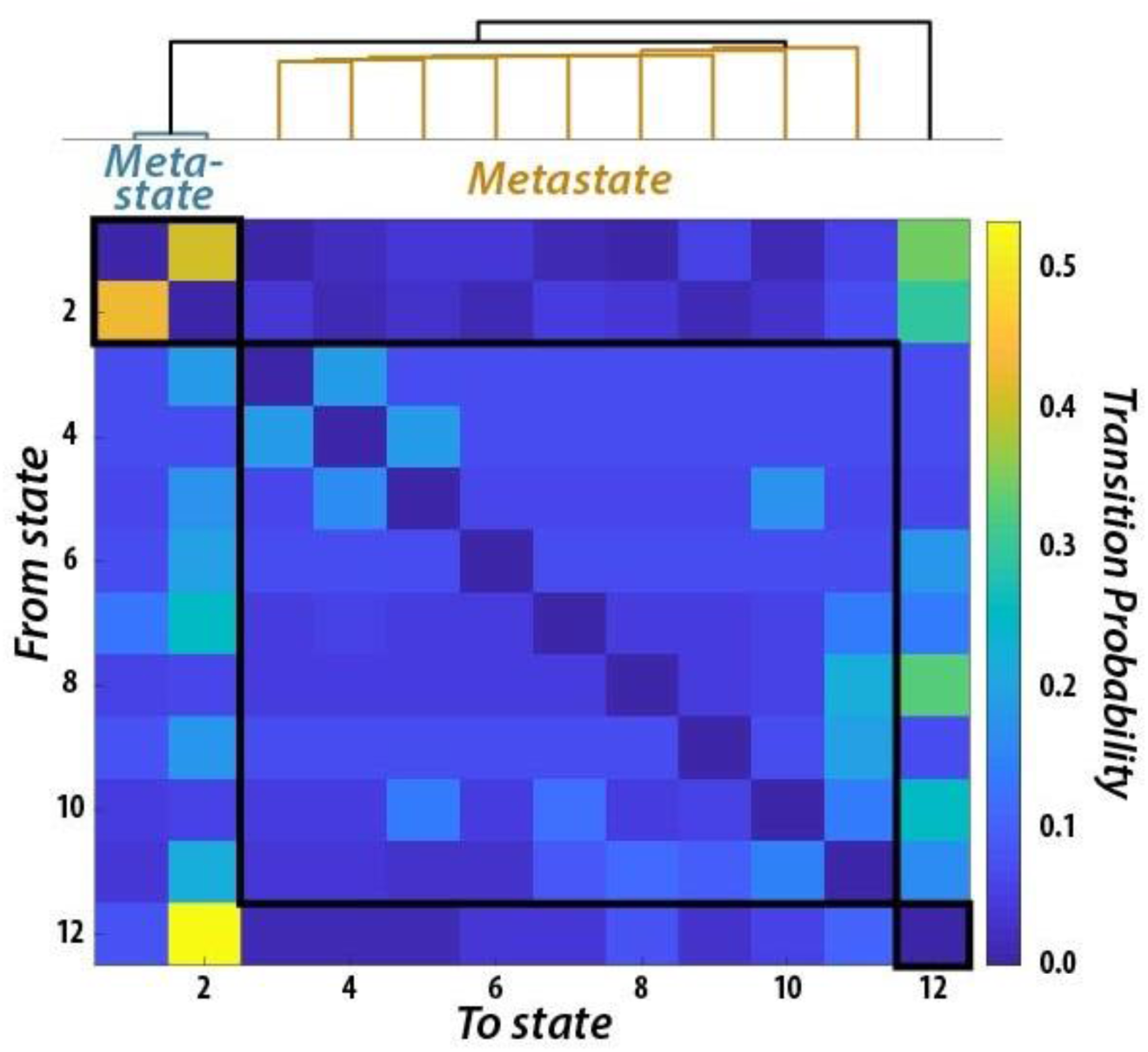
Metastates Resulting from the Temporal Clustering of Brain Dynamics. Probability, across all participants, of transitioning from one state to another. Clustering of the 12-state transition probability matrix revealed a temporal hierarchy wherein groups of states preferentially transitioned within groupings compared to across groupings. Two groupings contained multiple states (i.e. “metastates”), while one state was clustered only with itself.

Next, we used the fractional occupancy (i.e., time spent in each state) of each state and metastate, as well as the mean switching rate between states and metastates (n = 17 in total), as input into a CCA to determine the relationship between time-varying FC characteristics and the behavioral factors (n = 8, see Methods for description of selection and validation process). We found two significant CCA modes using the same permutation testing and cross-validation procedure as employed for static FC (Figure 4A).

**Figure 4.**
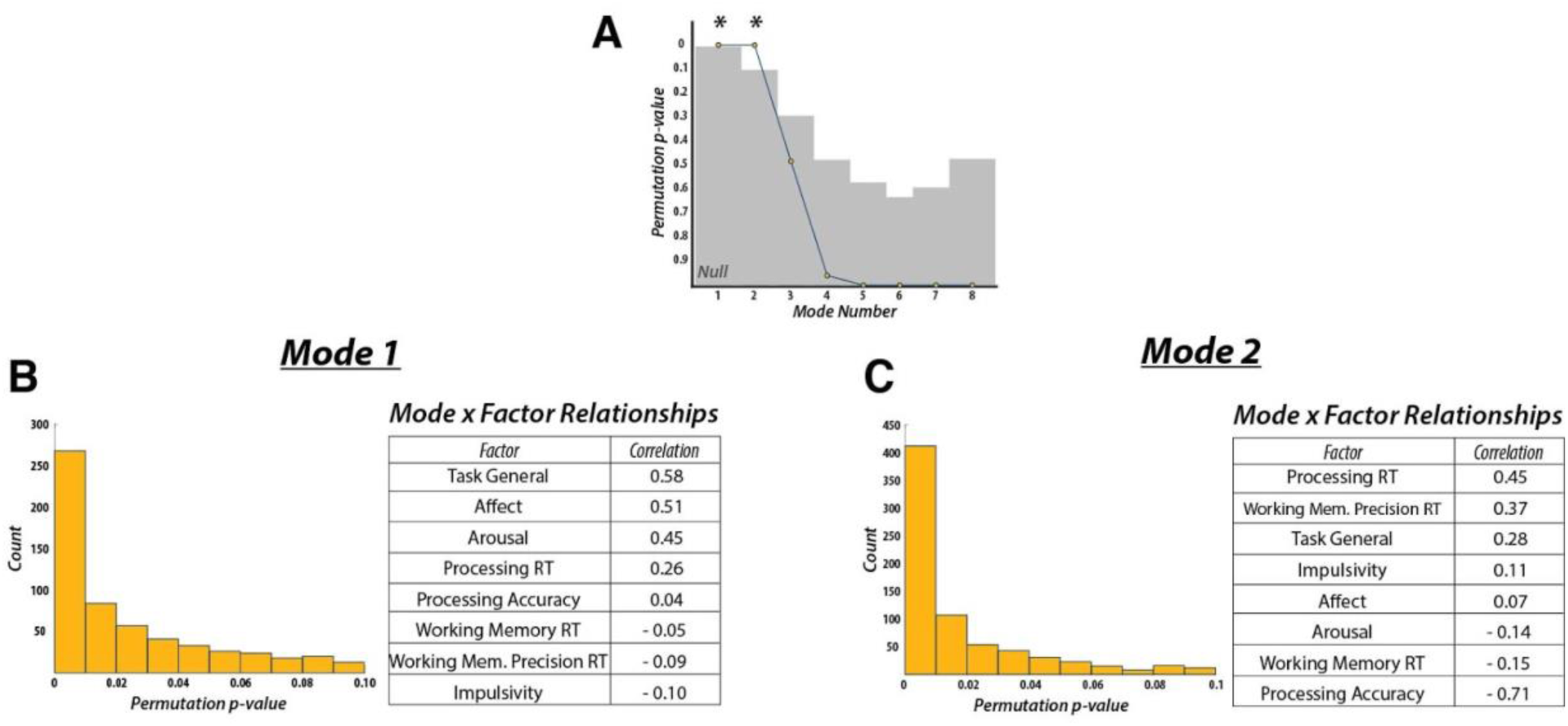
Canonical Correlation Analysis – Time-Varying Functional Connectivity. (A) CCA performed on measures of time-varying FC revealed two significant modes of covariation. Results of the cross-validation procedure and post-hoc correlations between (B) mode 1 and (C) mode 2 revealed that both modes were highly robust (assessed across 1,000 folds) and were sensitive to different sets of behavioral features. Whereas mode 1 largely matched the mode discovered with static measures of FC, mode 2 was instead sensitive to task- and trial-specific measures of behavior. RT: Reaction Time.

The first mode was defined by positive weights for “Task General” (r = 0.58), “Affect” (r = 0.51), “Arousal” (r = 0.45), and “Processing Reaction Time” (r = 0.26) factors, showing a similar pattern to the mode obtained from static FC (Figure 4B). Specifically, “Task General” and “Affect” loaded highest, while “Impulsivity” (r = −0.10) loaded most negatively (although its loading was greatly reduced compared to the previously discovered static mode). All other loadings fell below an absolute value of 0.09.

The second mode exhibited different behavioral weights when compared to the first time-varying mode. Here, “Task General” (r = 0.28), “Affect” (r = 0.07), and “Arousal” (r = −0.14) factors had substantially lower weights. Instead, “Processing Reaction Time” (r = 0.45) and “Working Memory Precision Reaction Time” (r = 0.37) factors loaded most highly on the positive end, while the “Processing Accuracy” (r = −0.71) factor loaded most negatively (Figure 4C). All remaining factors had weights below an absolute value of 0.15.

Similar to a previous analysis on the differentiable contributions of static and time-varying FC (Liégeois et al., 2019), we found that time-varying FC, while showing some similar relationships to behavior as static FC, could also distinguish relationships with more task-based measures of behavior. However, by using more specific measures of working memory (i.e. match-to-sample vs. free recall, accuracy vs. reaction time), task processing, and cognitive control, we were additionally able to determine that the second time-varying CCA mode distinguished unique behaviors associated with task performance. Specifically, the mode was defined by a separation (i.e., a positive-negative split in post-hoc correlations) between reaction time and accuracy, thus revealing within-task effects that previously had not been interrogated.

To further characterize each state obtained from the HMM, we overlapped their spatial profiles with those of canonical rsFC networks (Smith et al., 2009). Qualitatively, we found that the two-state metastate overlapped with two distinct task-positive networks (i.e., fronto-parietal and somatomotor networks; Figure 5). The nine-state metastate overlapped with a larger variety of networks, including the default mode, executive, and visual networks (Figure 5).

**Figure 5.**
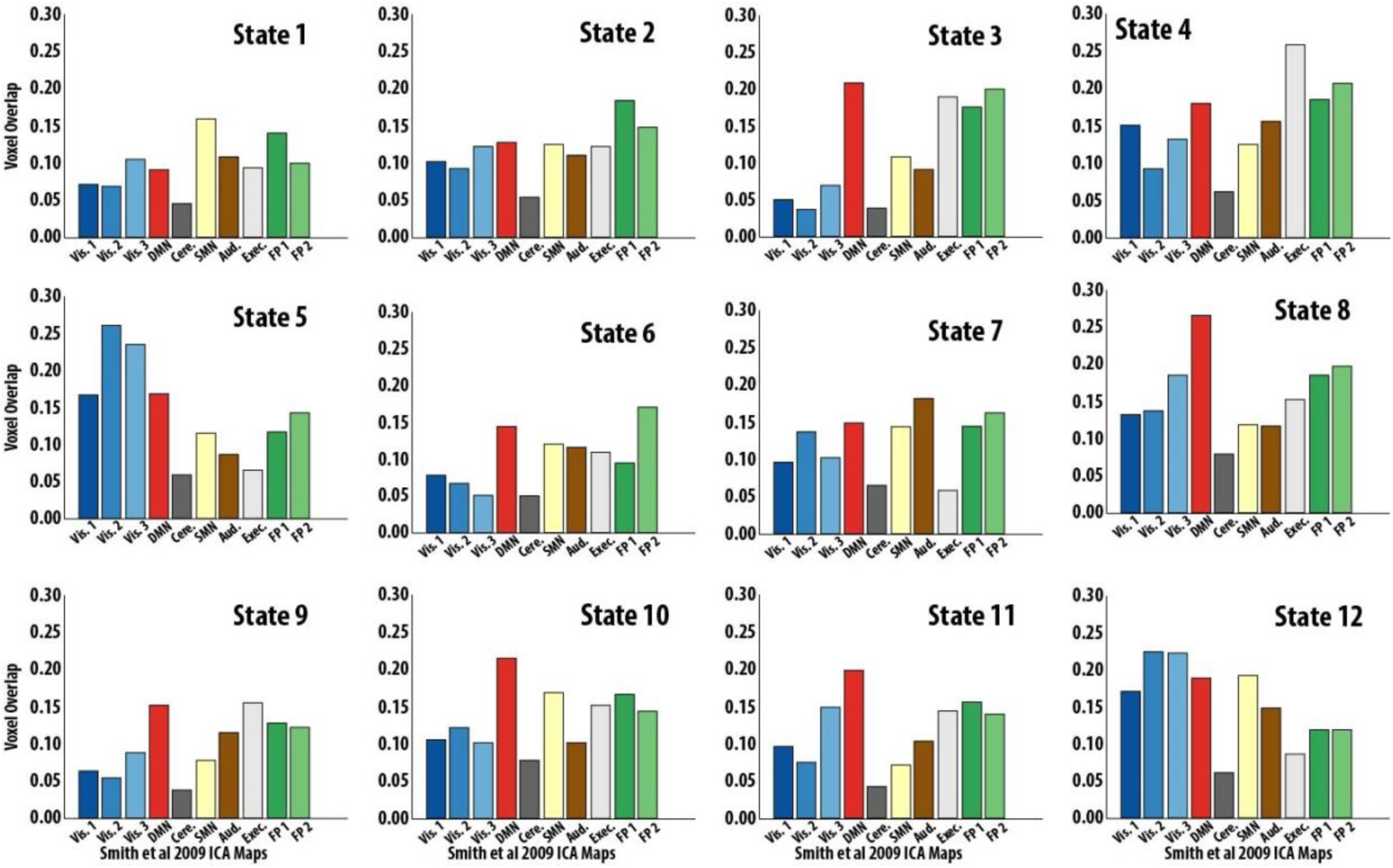
HMM State Activation Map Overlap with Resting-State Networks. Voxel overlap proportion for each HMM-derived state to the ten resting-state ICA maps from Smith and colleagues (2009). Ordering of states matches that of Figure 3. Specifically, states 1 and 2 clustered together in one metastate, states 3-11 in another metastate, and state 12 clustered alone.

## Discussion

Using CCA, we investigated the relationship between complex measures of human behavior and both static and time-varying rsFC in healthy young adults. Overall, we found a single CCA mode between the behavioral measures and static FC. In contrast, we found two CCA modes relating behavior and time-varying FC. Of these two modes, the first one resembled its static counterpart, while the other appeared to be distinct in that it was more sensitive to measures of task-specific behavioral variability. We thus argue that time-varying FC describes nuanced brain-behavior relationships distinctly from that which is captured by static FC.

Measures of static FC typically consider average FC over a prolonged period of time (e.g., several minutes of an fMRI scan) and have been used extensively to study the functional organization of the brain during rest and performance of a task (Cole et al., 2014, Cohen & D’Esposito, 2016). We first used nodal bidirectional FC edge strength quantified across the entire session in a CCA to relate the brain’s intrinsic static functional organization to behavior. The CCA revealed a significant relationship between these static measures and our behavioral factors. Measures of affect and impulsivity determined the main positive and negative directions of this mode, respectively. Additionally, and to a lesser extent, the positive direction of this mode was also characterized by an assortment of accuracy and response time behavioral measures derived from tasks sensitive to working memory (Spatial WM), cognitive control (Stroop), and processing speed (DSST). These results indicate that static FC likely tracks participant-level personality measures present during the scanning session (e.g., affective state). Unsurprisingly, static connectivity is also sensitive to measures of task performance that likely characterize stable behavioral features of the individual (i.e., general, multi-task performance, including working memory). As such, these results largely confirm the findings of previous studies on static rsFC’s predictive power in regard to certain measures of human behavior.

However, it has been shown that FC, including measures from resting-state protocols, is likely a dynamic process whereby fluctuations in regional connectivity occur rapidly (Lurie et al., 2019). Given the rate at which these fluctuations occur, one hypothesis regarding their cognitive relevance is that these fluctuations might better encode behavioral information reflecting ongoing cognitive demands, as compared to a general characteristic which would remain stable over the span of minutes, hours, or days. Previously, Casorso and colleagues (2019) assessed a similar, albeit broader, hypothesis by extracting time-varying rsFC components and submitting these to a CCA along with a selection of behavioral measures collected in the Human Connectome Project. Two modes of covariation were found between their time-varying connectivity components and measures of human behavior; however, no analysis of static FC was made against which to compare. One of the discovered modes was largely defined by positive post-hoc correlation values for measures of vocabulary comprehension and working memory, and negative values for measures tapping prosocial behaviors. The second mode was defined by positive post-hoc correlation values for measures tapping visuospatial orienting and emotional processing, and negative values for measures of inter- and intrapersonal processing and wellbeing. Although a critical step forward in the analysis and validation of time-varying FC’s relevance to human behavior, this study did not address the nature of how time-varying FC relates to behavior in a unique manner compared to static FC measures. Specifically, the behavioral measures used preclude the ability to measure processes that likely vary from trial to trial, as task-specific measures of reaction time and accuracy are, by design, combined to create composite scores across trials in the measures taken from the Human Connectome Project. In our experiment, we used behavioral measures that separately tracked processes related to stable (e.g., accuracy) vs. time-varying (e.g., reaction time) aspects of behavior to better assess our hypothesis.

Using measures of time-varying FC calculated from fitting a HMM to our rsFC data, we investigated whether CCA would reveal modes of population covariation sensitive to measures of behavioral variability. Our analysis resulted in two significant modes, whereby one mode largely resembled the mode discovered with static measures of FC. A comparison of this mode’s behavioral factor weights with its static counterpart suggests that these two modes might encode similar behavioral components. The primary difference between these modes is that this time-varying FC mode carried a highly positive weight for measures of drowsiness and fatigue, potentially reflecting a sensitivity of time-varying FC to neural and physiological correlates of arousal (Patanaik et al., 2018).

Whereas one of the time-varying modes reflected a largely similar, but not identical, behavioral profile as the static FC mode, the other time-varying mode reflected a more unique behavioral profile. High positive weights were associated with response time measures for tasks that assessed working memory and processing speed, while a strong negative weight was found for the measure of accuracy on the processing speed task. Characterized by measures of trial-by-trial response variability, this mode’s positive end potentially reflects a greater sensitivity to behavioral dynamics that occur on a more rapid timescale compared to what static FC is likely sensitive. In addition, the separation of measures of response variability and overall response accuracy, especially within the same task, reveals that time-varying FC is likely capable of disentangling unique behavioral components within the same task. Although our static FC mode did show some sensitivity to a measure that captures response variability, the distinction between stable and time-varying components of behavior was not present as is seen in our second time-varying mode. Overall, it is possible that this time-varying mode captures the relationship between brain dynamics and the measures of trial-by-trial behavioral variability within complex measures of human behavior.

The manner by which time-varying fluctuations in FC during the resting state relate to independent measures of human behavior remains unresolved. It is known that the spatial organization of functional connections changes in response to different tasks compared to rest (Cole et al., 2014; Cohen & D’Esposito, 2016). Specifically, inter-network connectivity is more predominant during tasks that require flexible cognition (i.e., working memory) compared to more rudimentary tasks such as executing specific finger-tapping sequences. Moreover, a previous report found that measures of global network integration and within-network connectivity (i.e., participation coefficient and module degree, respectively), when assessed in a time-varying manner, varied throughout the performance of tasks and tracked the cognitive complexity of the current task demands (Shine et al., 2016). Thus, one hypothesis as to how resting dynamics relate to behavior is that the dynamic interactions within and between these networks observed during tasks can be recapitulated during periods of wakeful rest. However, it should be noted that the dynamic interactions that occur during task performance are likely more constrained than during rest due to the confined cognitive context required by task performance. Resting-state dynamics can serve as a “baseline” repertoire that can potentially index the extent to which FC reconfigures during task and, in turn, track behavioral performance (Liégeois et al., 2019). Although the current report did not collect task-based fMRI data, it will be crucial for future studies on the behavioral relevance of time-varying FC to assess this possibility.

It is also important to emphasize the spatio-temporal signature of these time-varying network interactions and what it means for behavioral performance. Methods such as the HMM investigate brain dynamics with high temporal resolution, thus extending previous methods showing reconfiguration of connectivity between different task blocks (Cohen, 2018). For example, Vidaurre and colleagues used a HMM to show how a motor task drives reconfiguration of large-scale networks on a timepoint-by-timepoint basis showing that task execution happens at faster timescales that had been previously undetected when interrogated using sliding window methods (Vidaurre et al., 2018). Regarding the spatial profile of the current HMM states, a visual and quantitative assessment of their overlap with canonical rsFC networks (Smith et al., 2009) suggested that our metastates had distinct spatial profiles. We identified a nine-state metastate spanning multiple networks including fronto-parietal, executive, default-mode, and visual networks. Integration of the “task-positive” and “task-negative” networks has been observed during motor tapping and autobiographical planning, suggesting a more mutually compatible role than previously believed (Fox et al., 2005), one that can facilitate goal-directed cognition (Spreng et al., 2010; Braga et al., 2013; Vatansever et al., 2015). On the other hand, the two-state metastate we identified, characterized by a more constrained spatial profile of fronto-parietal and somatomotor networks, potentially reflects networks specific to task execution. The differences in spatial topography of the two-state versus nine-state metastates may provide insight regarding the different behavioral relationships we found with static versus time-varying FC. The flexible interaction of activity across each metastate’s respective individual states might allow for the encoding of information to which static measures are insensitive. Although static measures are capable of reflecting multi-network interactions, they are incapable of tapping into the specific temporal patterns through which these network interactions occur. Further investigation of the spatial patterns of these states is needed.

In conclusion, the current study demonstrates that static and time-varying FC are differentially associated with behavior. We argue that via integration across multiple networks at different temporal scales, time-varying FC is associated with both trial-by-trial and stable behavioral measures, while static FC is associated with participant-level personality measures and measures of stable task-general performance. These results demonstrate that it is important for future studies to look at both the static and temporal aspects of FC to more fully delineate the behavioral contributions of each.

## Supplementary Materials

**Figure S1.**
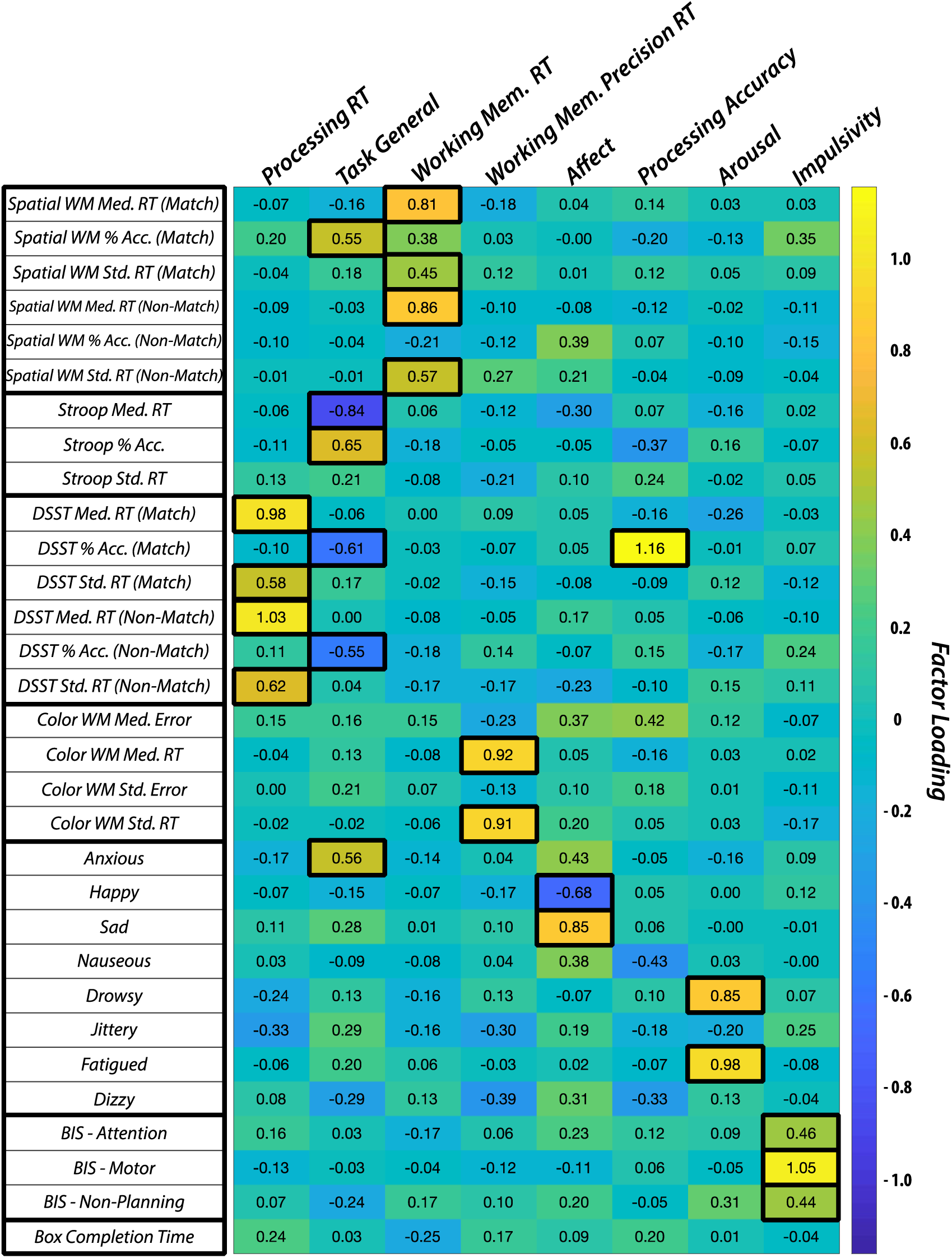
Factor Loadings in Behavioral Data. Shown are the factor loadings across the 8 derived factors for each of the 31 behavioral measures. To aid interpretability, certain values are highlighted to show the clustering of loadings within groups. Factors were assigned names based on the behavioral variables that loaded most heavily on them. RT: Reaction Time.

**Figure S2.**
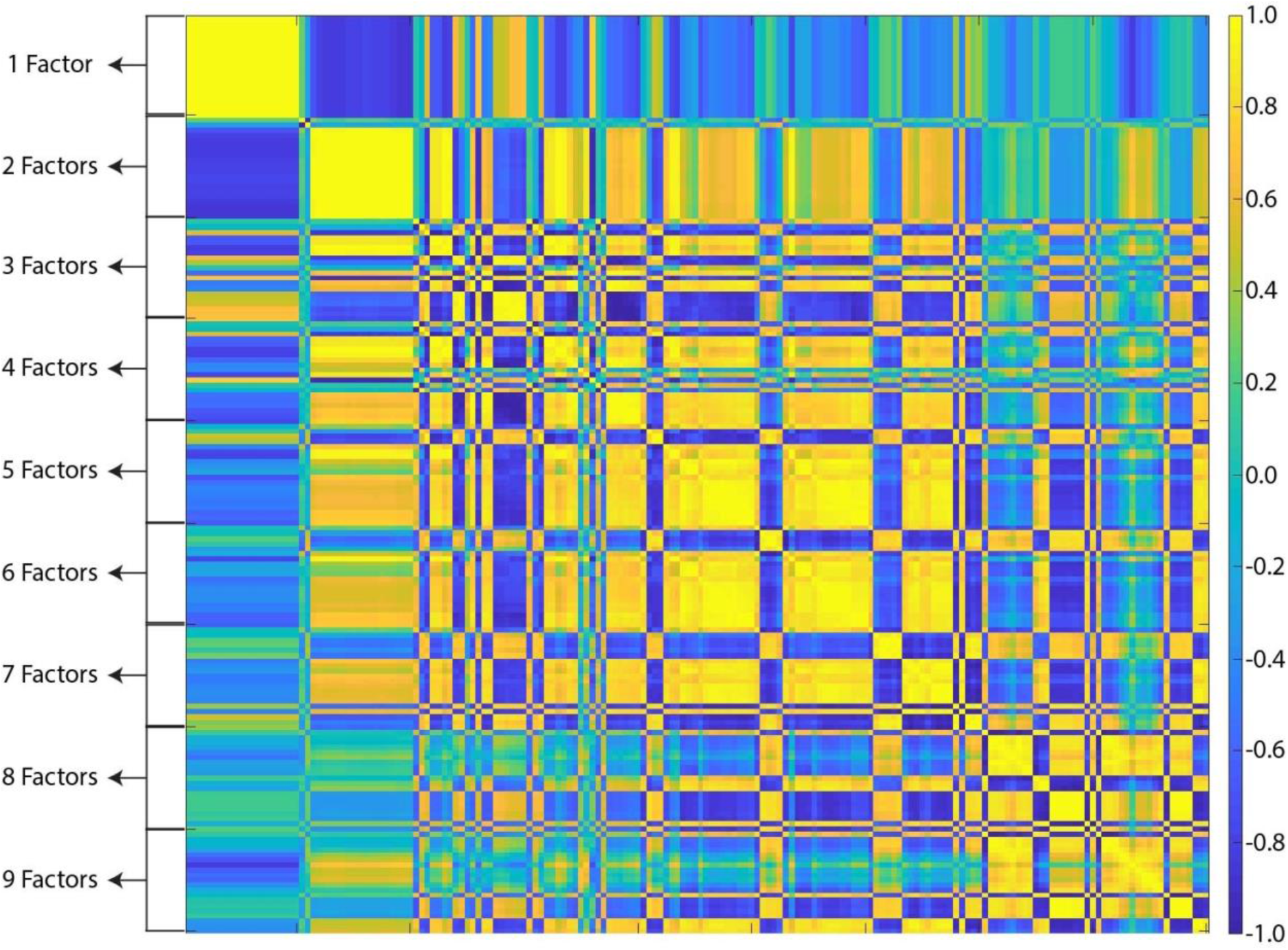
Validation of Static Functional Connectivity CCA. Similarity between modes of covariation are plotted across varying numbers of behavioral factors (1-9) and edge strength principle components (1-20). Similarity was measured as the Pearson correlation coefficient between each mode’s vector of 31 behavioral post-hoc correlations. The plot is organized such that for each CCA run, connectivity data in varying principal component space is grouped together within chunks of behavioral data in a fixed factor-space. For example, the first row represents CCA being run on behavioral data in 1-factor space and connectivity data in 1-PC space; the 10th row represents CCA being run on behavioral data in 1-factor space and connectivity data in 10-PC space; the 15th row represents CCA being run on behavioral data in 2-factor space and connectivity data in 5-PC space, etc. Essentially, only one mode is found across all PCs when in 1- and 2-factor space, with the discovered modes being identical within factor space, and almost exact inverses of each other across factor spaces. Modes discovered from CCA run on data in 3-factor space and above, across all PCs, almost entirely represent the same relationship as each other.

**Figure S3.**
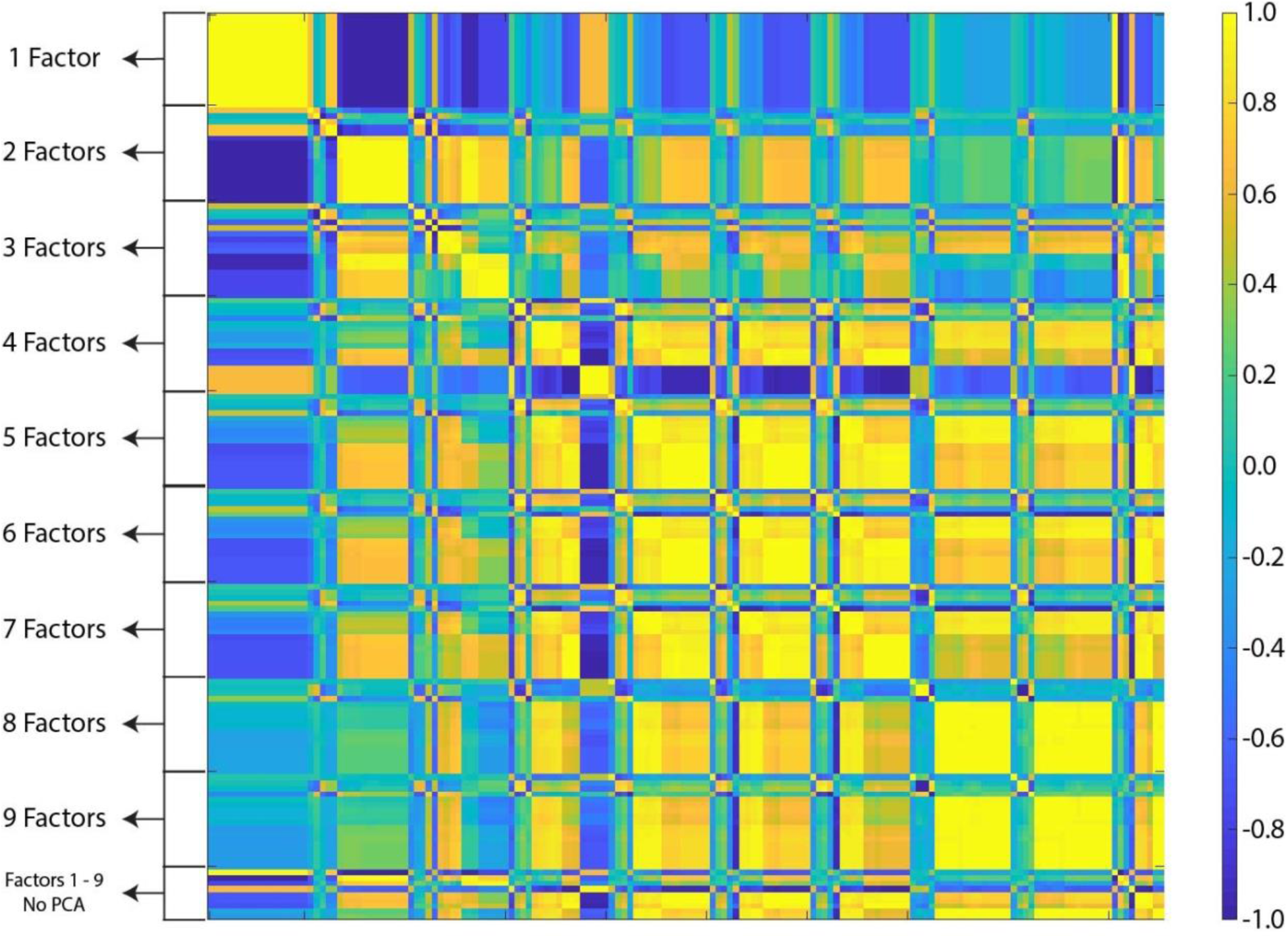
Validation of Time-varying Functional Connectivity CCA. Similarity between modes of covariation are plotted across varying numbers of behavioral factors (1-9) and edge strength principle components (1-17, capped by number of HMM variables included). Similarity was measured in the same way as for the static FC CCA validation procedure using Pearson correlation. Unlike in the static FC case, we additionally assessed whether the discovered modes differed as a function of whether PCA was, or was not, run on the HMM data. The final 9 rows represent the modes discovered from CCA run on behavioral data in 1-through 9-factor space when PCA was not performed on the HMM data. As can be seen, the modes discovered when CCA was run on the raw HHM data almost perfectly match those found when PCA was run on the HMM data. A corresponding plot for post-hoc correlations computed from modes greater than 1 is inappropriate as we had no *a priori* knowledge that more than 1 significant mode of covariation would be discovered, and thus only the first mode of covariation can be compared across datasets.

